# Identification of a specific set of genes predicting obesity before phenotype appearance

**DOI:** 10.1101/2025.02.28.640741

**Authors:** Céline Jousse, Laurent Parry, Gwendal Cueff, Marion Brandolini-Bunlon, Jérémy Tournayre, Alain Bruhat, Anne-Catherine Maurin, Cyrielle Vituret, Julien Averous, Yuki Muranishi, Pierre Fafournoux

## Abstract

Obesity poses significant health and socioeconomic challenges, necessitating early detection of predisposition for effective personalized prevention. To identify candidate predictive markers, our study used two mouse models: one exhibiting interindividual variability in obesity predisposition and another inducing metabolic phenotypes through maternal nutritional stresses. In both cases, predisposition was assessed by challenging mice with a high-fat diet. Using multivariate analyses of transcriptomic data from white adipose tissue we identified a set of genes whose expression correlates with an elevated susceptibility to obesity. Importantly, the expression of these genes was impacted prior to the appearance of any symptoms. A prediction model, incorporating both mouse and publicly available human datasets, confirmed the discriminahve capacihes of our set of genes across species, sexes, and adipose hssue deposits. These genes are promising candidates to serve as diagnostic tools for identifying individuals at risk of obesity.

## INTRODUCTION

In developed countries, metabolic diseases – including Type II Diabetes (T2D) and obesity – have emerged as significant health concerns, causing disability and imposing substantial economic and social burdens on societies. These diseases encompass various conditions that may share common underlying causes and often manifest as syndromes, of which metabolic syndrome (insulin resistance/T2D, obesity, dyslipidemia, hypertension) is a prominent example. Metabolic diseases typically develop over time, often progressing silently for many years before clinical symptoms appear. Individuals have a variable susceptibility to metabolic diseases. The speed and extent of metabolic changes leading to disease onset are influenced by both intrinsic and environmental factors. Detecting predisposition to metabolic dysfunction before clinical symptoms emerge could contribute to prevention and the development of personalized treatments, leading to more effective personalized medicine.

Metabolic diseases have diverse causes, combining genetic, epigenetic, and environmental factors. Genetic factors have provided insights into the pathogenesis of T2D, but the genetic variants identified are estimated to account for only a small proportion (approximately 5 to 10%) of the heritability of this disease ^1^. It is now widely recognized that epigenetic factors play a crucial role in determining an individual’s health status over their lifespan. In particular, the perinatal period represents a critical window for such modifications. Indeed, a maternal nutritional mismatch during pregnancy and/or in the early weeks/months after birth significantly contributes to the development of obesity, metabolic syndrome, and diabetes ^2^.The fetus responds to maternal nutritional stresses through specific adaptations at the cellular and molecular levels, which permanently affect its physiology and metabolism, persisting even when the initial stress has been resolved. Thus, early environmental stressors affect the epigenome and subsequently elicit sustained responses, modifying gene expression patterns and phenotypes in adulthood. Nutritional programming has been clearly demonstrated in both animals and humans, leading to the emergence of the “Developmental Origins of Health and Disease” (DOHaD) hypothesis ^3–8^.

We propose genes expression patterns in adipose tissue that are influenced by perinatal imprinting could serve as early predictive markers of a predisposition to metabolic disease development. To identify potential early predictive biomarkers of at-risk populations, we established two complementary animal models: (i) a large cohort of inbred mice to study interindividual variability in terms of predisposition to obesity and (ii) an experimental model investigating the effects of maternal nutritional stress and its timing (during gestation and/or lactation) on the metabolic phenotype of adult offspring. In both cases, to determine the predisposition to obesity, we exposed mice to a high-fat diet (HFD) challenge. Adipose tissue biopsies were taken from each animal before and after the HFD-challenge. This enabled us to analyze gene expression patterns prior to the appearance of the obesity phenotype, thus capturing early molecular indicators of predisposition to obesity.

By analyzing gene expression in both experimental models, we identified a set of genes that correlate with an elevated susceptibility to obesity and whose expression was impacted prior to the emergence of symptoms. We compared our results to results for human subjects using publicly accessible human expression datasets, and used prediction modeling to show that these genes have discriminative capacities whatever the specie (either human or mouse), the sex and the localization of the white adipose tissue depot. The genes identified may hold potential for the development of innovative diagnostic tools to identify individuals who are at risk of developing metabolic diseases. They could also be used to monitor interventions and treatment efficacity leading to more effective management of metabolic diseases.

## RESULTS

### Innate proneness to diet-induced obesity (DIO) is highly variable between individuals

In any given population, whether human or mouse, there are significant variations in susceptibility to the development of obesity and metabolic disorders, particularly in response to aging and/or dietary factors (high fat and sugar) ^9,10^. In a first set of experiments, we assessed the weight gain in responses to a nutritional challenge (HFD) of a cohort (n=27) of inbred (i.e., genetically very close) adult mice presenting a uniform basal phenotype (as depicted in Figure 1A). As shown in Figure 1B, at timepoint T0, the distribution of body weight and fat mass among the mice followed a normal distribution. However, after the 18-week HFD-challenge (T18), a wide range of sensitivities to diet-induced obesity (DIO) were observed, with the emergence of three distinct groups. Specifically, certain mice exhibited minimal weight gain and were thus classified as DIO-resistant, whereas others exhibited excessive weight gain and were considered prone to DIO. An intermediate group with intermediate weight gain was also identified.

**Figure 1:**
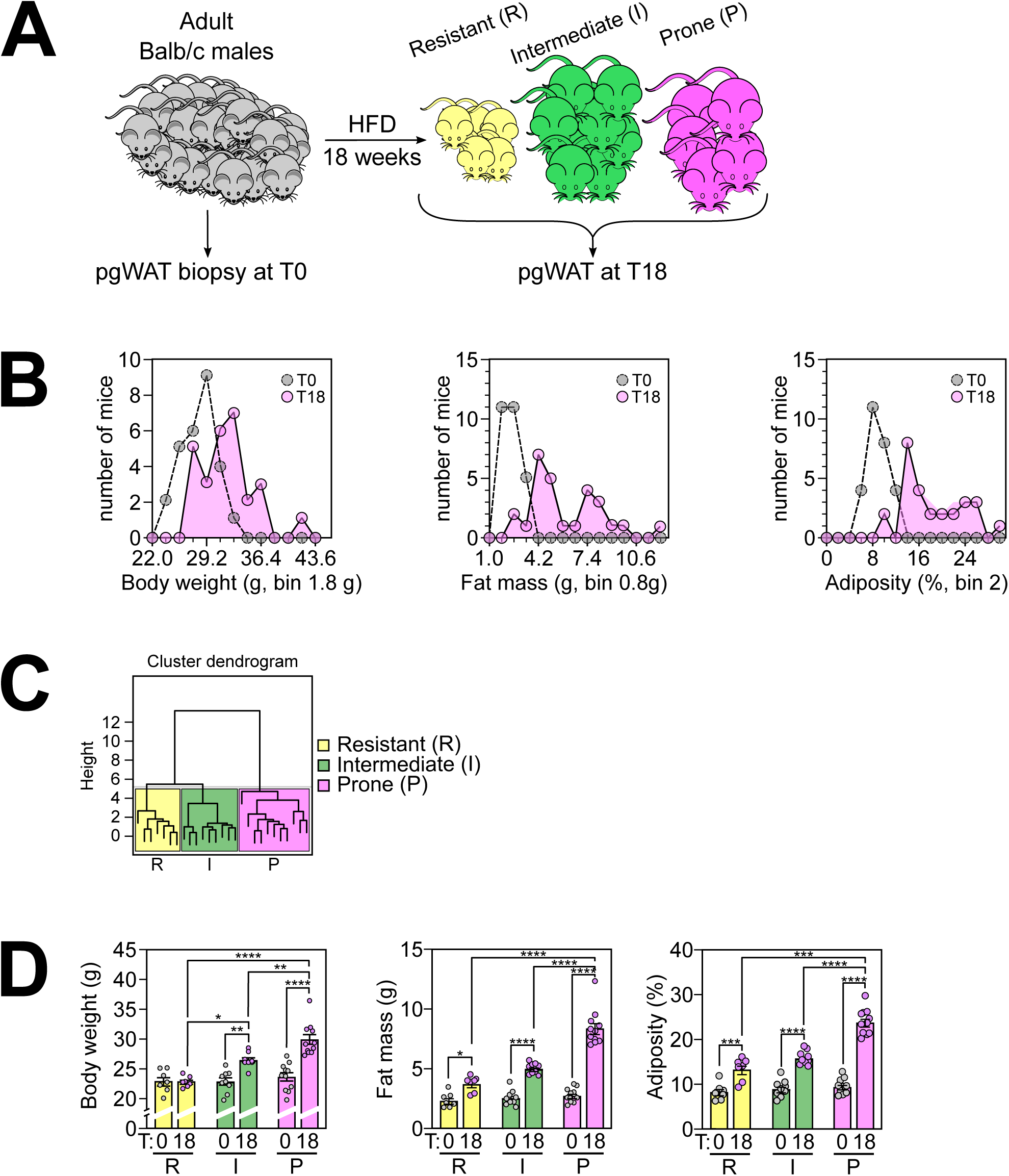
Innate proneness to diet-induced obesity (DIO) exhibits a large degree of variability among individuals. **A: Experimental model.** 4-month-old Balb-c males mice were monitored (n=27). At T0, perigonadal WAT (pg-WAT) biopsies were taken. Animals were allowed to recover for 1 week. Subsequently, they were fed an experimental high-fat diet (HFD) for 18 weeks after which they were sacrificed (T18) to harvest perigonadal WAT. Schematicaly, at the end of the HFD challenge, the mice exhibit different sensitivities to obesity and will thus be categorized into 3 groups: R mice are resistant to Diet Induced Obesity DIO, I mice have an intermediate phenotype, and P mice are prone to DIO. **B: Phenotypic characterization of mice based on sensitivity to DIO.** Individual parameters such as body weight (g), fat mass (g), and adiposity (%) were measured at T0 (before HFD-challenge) and at T18 (after 18 weeks consuming a HFD). Graphs show the distribution of the number of mice according to each parameter. Gray dots represent the animal before HFD-challenge, filled dots represent animals after HFD-challenge (T18). In each case, three groups of mice exhibiting different behavior regarding diet-induced changes in physical parameters are clearly identified. **C: Hierarchical Clustering Analysis (HCA)** considers all the physical parameters measured (i.e. body weight, fat mass, and adiposity at T18, delta% body weight, delta% fat mass, and delta% adiposity) to classify mice in three groups: R mice are resistant to DIO (n=7), P mice are prone to DIO (n=11), and I mice exhibit an intermediate phenotype (n=9). **D: Body weight, fat mass and adiposity before and after HFD-challenge for each group R, I and P.** Body weight (g), fat mass (g), and adiposity (%) at T0 (gray dots) and T18 (pink dots) for R, I, and P groups (n= 7, 9, 11 respectively). Bars represent the mean for each group and error bars correspond to SEM. One-way ANOVA p-value is indicated as follow: * p ≤ 0.05; ** p ≤ 0.01; *** p ≤ 0.001; **** p ≤ 0.0001.

To comprehensively consider all physical parameters including body weight, fat mass, and adiposity at T18, delta% body weight, delta% fat mass, and delta% adiposity, a Hierarchical Clustering Analysis (HCA) was conducted (Figure 1C). The results confirmed that three distinct groups of mice were present following the HFD-challenge. Based on this *a posteriori* HCA (Figure 1D), for the first group, referred to as Resistant (R), body weights at T0 and T18 were not significantly different, with only slight increases in fat mass and adiposity. The second group, referred to as Intermediate (I), exhibited a significant increase in all three parameters between T0 and T18. The extent of the increase was higher than observed for the R group. Finally, the third group, referred to as Prone (P), gained considerable body weight, fat mass, and adiposity, to significantly higher extents than the other groups. These animals were therefore considered highly susceptible to DIO. Our results confirm that, within an ostensibly homogeneous population, some individuals are highly sensitive to DIO and are therefore prone to developing associated pathologies, whereas others are more resistant.

These observations indicated that prior to the HFD-challenge, at T0, none of the parameters measured (body weight, fat mass, and adiposity) could reliably predict an individual’s response to a HFD. This conclusion was further supported by attempts to develop a predictive model using partial least squares discriminant analysis (PLS-DA) based solely on the physical parameters measured at T0, as demonstrated through a permutation test (Figure S1A, left panel). In contrast, PLS-DA and permutation tests conducted on the T18 data (Figure S1A, right panel) confirmed that the three groups identified are indeed significantly different and can be discriminated based on their physical parameters.

### Identification of a set of genes as early predictive markers of predisposition to DIO

Our main objective was to identify genes with an expression pattern correlating with predisposition to DIO. We specifically focused our study on perigonadal white adipose tissue (pg-WAT) due to its metabolic relevance and its ease of accessibility for biopsies. One week before the change in diet (T0) a pg-WAT biopsy was taken. At the final time point (T18) following the 18-week HFD-challenge, pg-WAT was taken at sacrifice.

Based on the clustering shown in Figure 1C, the pg-WAT gene expression profiles were compared for the Resistant, Prone, and Intermediate groups at T0 and T18. The workflow used for this analysis is illustrated in Figure 2A. First, differential expression analysis was conducted on the microarray gene expression data (Array Express E-MTAB-13877 and E-MTAB-13878) using the Limma package. A total of 2356 mRNA species were differentially expressed at T0, and 1020 mRNA species were differentially expressed at T18 according to the group factor (I, R, or P). This preliminary statistical analysis was later used to refine the gene list. Second, PLS-DA was performed on the same dataset to identify variables that could effectively identify individuals belonging to the three groups at each time point. To assess the reliability of the predictive models obtained and ensure they were not affected by overfitting or excessive randomness, permutation tests were conducted by randomly permuting (500 times) the individuals in the initial dataset (Figure S1B). The error rates obtained with all the permuted data were higher than with the original dataset, thus the validity of the PLS-DA models obtained was confirmed. To assess the contribution of each variable to the predictive models, Variable of Importance in Projection (VIP) scores were calculated. A threshold VIP of ≥ 1.5 was applied, as it represents a robust criterion to select variables of interest. Following application of this threshold, we retained 3387 mRNA species at T0 and 1746 mRNA species at T18 as potential variables of interest in the analysis (Figure 2B).

**Figure 2:**
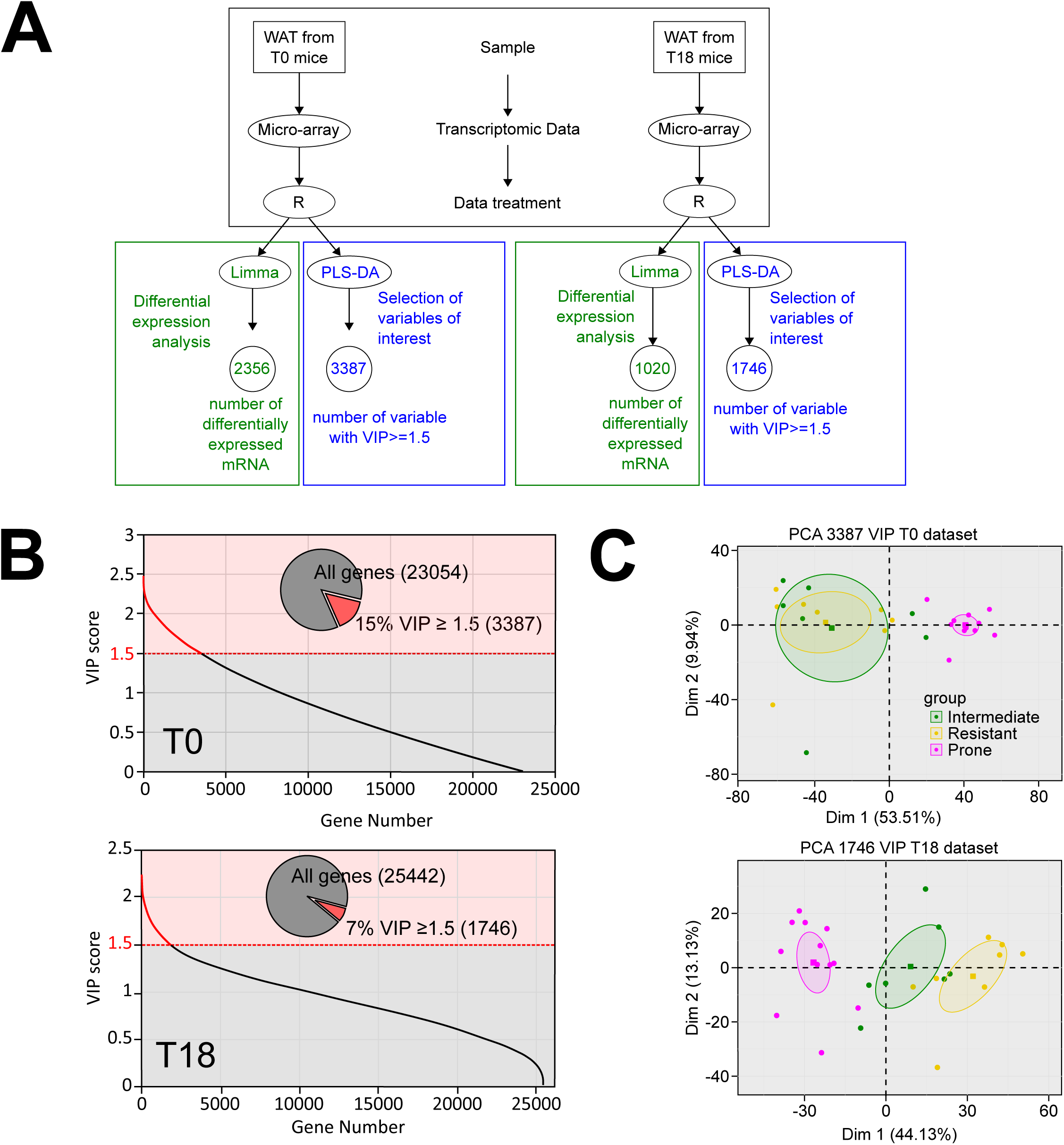
Identification of a set of genes as early predictive markers of predisposition to DIO. **A: Workflow to select genes of interest.** Micro-arrays were performed on RNA extracted from pg-WAT harvested at T0 and T18 from mice in the three groups R (n= 8), I (n= 7), P (n= 12). Raw data were treated with the R package Limma prior to differential expression analysis (green part of the workflow). The mixOmics R package was used for PLS-DA (blue part of the workflow) in order to identify important genes based on their VIP scores. A VIP score is a measure of a variable’s importance in the PLS-DA model. It summarizes the contribution of a variable makes to the model. **B: Variable of importance in projection (VIP) scores for each gene used in the PLS-DA.** Genes contributing meaningfully to the PLS-DA model with a VIP score >1.5 constitute respectively 15% and 7% of the genes tested for T0 and T18 data (inset pie-chart). The y-axis corresponds to the VIP scores for each variable on the X-axis. Red part of the curve corresponds to genes with the highest VIP scores (>=1.5) and thus are the most contributory genes in class discrimination in the PLS-DA model. **C: Principal Component Analysis (PCA).** PCA was performed using the set of 1746 mRNA selected by PLS-DA at T18 and 3387 mRNA selected by PLS-DA at T0. Data from groups I (green dots), R (yellow dots), and P (pink dots) are plotted, along the first (X) and second (Y) principal components. Ellipses represent the 95% confidence interval and squares represent the barycenter of each group.

Principal Component Analysis (PCA) was then applied to these data – the set of 3387 mRNA species at T0 and 1746 mRNA species at T18. The aim was to evaluate the ability of these mRNA species to effectively classified mice into their original group assignments. The results of the PCA analysis, as shown in Figure 2C, demonstrated that these selected mRNA species had a significant discriminatory capacity, particularly for mice in group P. Thus, these mRNA species could be useful indicators to identify individuals with a predisposition to DIO.

### The metabolic fate of offspring is primarily determined by the mother’s nutritional status

Considering that in the experimental model depicted above, the animals are genetically very close, the origine of the differences observed in susceptibility to obesity could potentially be attributed to epigenetic factors. In line with the principles of the Developmental Origins of Health and Disease (DOHaD) concept, susceptibility to obesity could be influenced by challenges experienced during perinatal life, especially depending on the mother’s nutritional status. In previous studies ^11,12^, we reported that mothers fed a Low Protein Diet (LPD) during gestation and lactation give birth to animals with increased basal energy expenditure and protection against high-fat-DIO. However, other groups reported that maternal undernutrition restricted to the gestational period leads to an opposite phenotype in offspring, making them prone to metabolic diseases and more sensitive to DIO ^13–17^. To further investigate this issue, we assessed the effects of the type and timing of maternal nutritional stress on predisposition to DIO among offspring. As shown in Figure 3A, a control group of males born to mothers fed a control diet (F1-CD, group A) was compared to the following experimental groups: males born to mothers fed a LPD during both gestation and lactation (F1-LPD, group B), males born to mothers fed a LPD during gestation only (F1-LPD-CD, group C), males born to mothers fed a HFD during lactation (F1-CD-HFD, group D), and males born to mothers fed a LPD during lactation only (F1-CD-LPD, group E). After weaning, all male offspring from these groups were fed a control diet. Various biological parameters were measured in adult male offspring from the five groups. Figure 3B (top-left) shows the body weight measured from 10 days and throughout life, up to 16 months. At 5 months, the weight of the animals is presented Figure 3B (bottom). This analysis reveals that males from group C showed no difference in body weight compared to the control group A, whereas males from groups B and E had lower body weight, and males from group D had higher body weight. Similar trends were observed when considering the fat mass of the animals (Figure 3B-bottom). It is interesting to note that maternal nutritional stress had a greater impact on fat mass than on lean mass, although both were affected. Representative photographs of animals from extreme groups at 10 days and 2 months of age further illustrate the considerable differences in body weight linked to maternal nutritional status (Figure 3B, top-right). Overall, these findings highlight the major impact of the type and timing of maternal nutritional stress during the perinatal period on the developmental trajectory of offspring.

**Figure 3:**
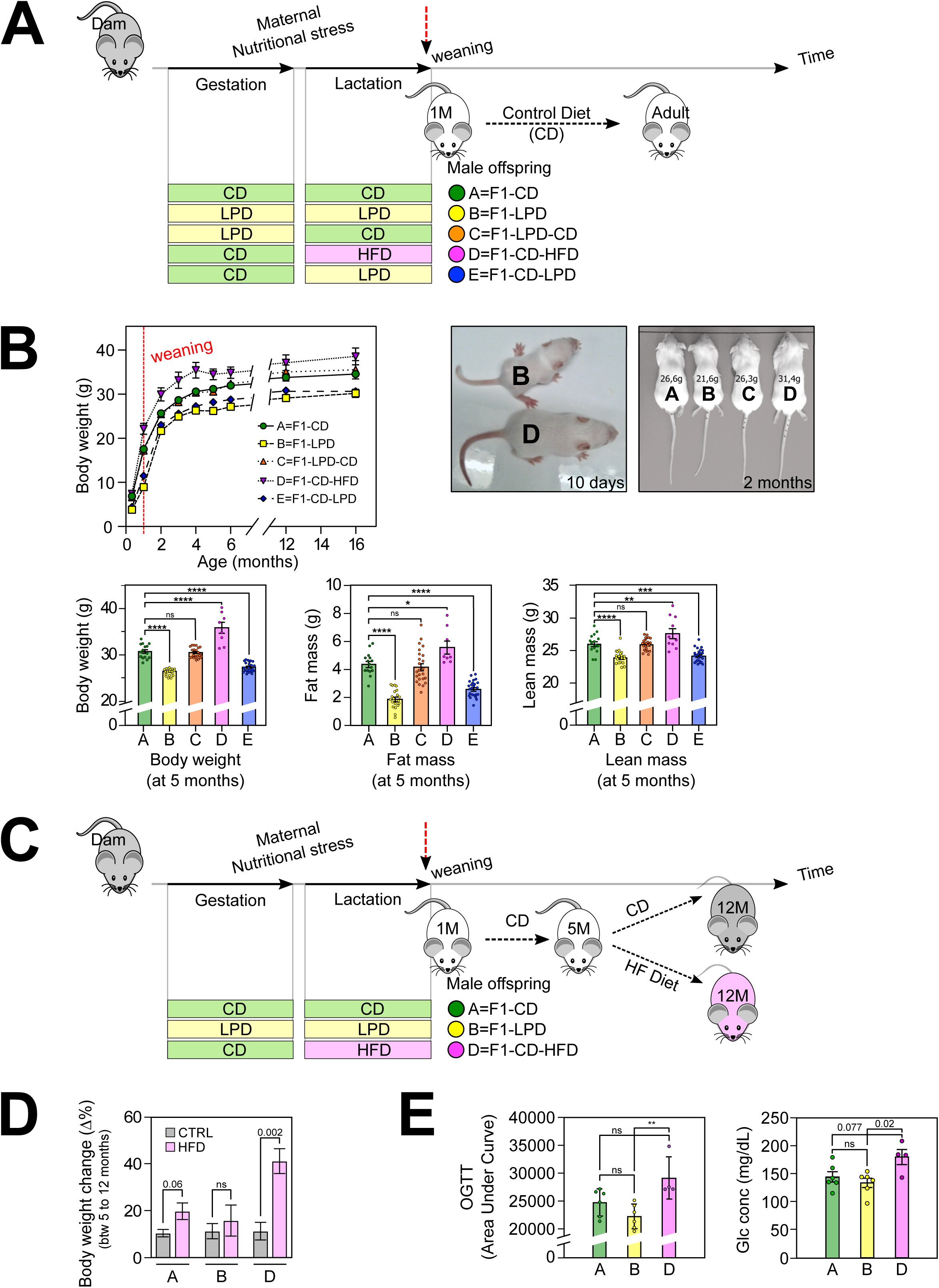
The metabolic fate of the offspring is primarily determined by the nutritional status of the mother. **A: Experimental model.** Two-month-old virgin BALB/c female mice fed a A03 chow diet were mated with BALB/c males. Gestating animals were isolated when a vaginal plug was detected, and fed the experimental diet as indicated. LPD and CD are isocaloric. At parturition, dams and litters were fed with the experimental diets indicated. Litters of different sizes were obtained from each group of pregnant female. Since the litter size is important in the offspring life trajectory, we considered only litters that have a total number of pups comprised between 4 and 10 to avoid extreme litter size. After weaning, male offspring from each group were housed individually with free access to CD. Moreover, to obviate any litter effects, animals used for each experiment were randomly chosen from different litters, and only a limited number of animals (n = 1 to 2) was used from each litter. **B: Phenotypic characterization of male offspring born from dams fed various diets during gestation and lactation:** (upper left panel) Male offspring born from dams fed experimental diets during gestation and lactation were weighed at Post-Natal Day 10 (PND10), every month from 1 to 6 months, and at 12 and 16 months. The results presented are the average of 3 independent experiments. (upper right panel) Representative mice from groups B and D at PND10 and from groups A, B, C, and D at 2 months. Body weight is indicated for each mouse. (bottom panel) Body composition parameters of 5-month-old A, B, C, D, and E male mice. Body weight, fat mass and lean mass are indicated in grams. All values correspond to mean +/- SEM for at least n=8 /group. One-way ANOVA p-value is indicated as follow: * p ≤ 0.05; ** p ≤ 0.01; *** p ≤ 0.001; **** p ≤ 0.0001. **C: Experimental model for the HFD-challenge on groups A, B, and D:** 5-month-old male offspring from groups A, B, and D were fed either a CD or a HFD from the age of 5 to 12 months (7-month HFD-challenge). **D: Body weight gain during HFD-challenge:** Body weight gain between the beginning (5 months) and the end (12 months) of the HFD-challenge was calculated and expressed as the weight gain relative to the initial weight, given in %. n>=4/group, One-way ANOVA p-value is indicated. **E: OGTT measured after HFD-challenge:** 12-month-old male mice from groups A, B, or D fed a HFD for 5 months were subjected to an Oral Glucose tolerance test. Area Under curve and starved glucose concentration were measured. n>=4, One-way ANOVA p-value is indicated is indicatedas follow: ns=not significant, ** p<0.01.

In subsequent investigations, we specifically focused on the control group (A) and the two experimental groups that displayed the highest and lowest body weight – groups D and B, respectively. Key metabolic parameters measured in young adults revealed that blood glucose levels and triglyceride levels were slightly lower in group B compared to group A, whereas triglyceride levels were slightly higher in group D (Figure S2). However, overall, the basic biochemical parameters such as glucose, triglycerides, and cholesterol varied little between groups, and remained within the range of physiological values in all cases. These findings indicate that, regardless of the experimental group, the animals showed no symptoms of illness.

To determine the predisposition of the offspring from each group toward obesity, young adult F1 animals (5 months old) were fed a HFD for 7 months, according to the experimental protocol presented in Figure 3C. As shown Figure 3D, control animals (group A) gained two-fold more weight when fed a HFD compared to a control diet (medium effect size; Cohen’s d = 0.54). In contrast, group D animals gained 4 times more weight when fed a HFD compared to a control diet (large effect size; Cohen’s d = 0.92). Interestingly, group B animals appeared to be protected against HFD-induced obesity, gaining only 1.5 times more weight when fed a HFD rather than a control diet. Figure S3A provides individual weight data for each mouse before and after the challenge period, either on a control diet or on a HFD. The data presented in Figure S3B also demonstrate that, in response to the HFD-challenge, changes in fat mass mirror the pattern observed for total body weight.

An oral glucose tolerance test (OGTT) was conducted at the end of the 7-month HFD- challenge. As shown in Figure 3E, the area under the curve for group D animals was the highest among the groups, suggesting a tendency toward insulin resistance. No differences were observed between the groups before the HFD-challenge (data not shown). Moreover, blood glucose levels were consistently higher in group D animals.

In summary, these findings demonstrate that group B animals are resistant to DIO, whereas group D animals are prone to DIO, indicating that maternal nutritional stress can produce a metabolic imprint in the offspring that significantly influences their susceptibility to metabolic disorders. Based on these results, groups B and D could be seen as equivalent to groups R and P identified in the previous experimental model, but obtained by induction rather than stochastically. These results reinforce the role of perinatal life as a major determinant in an individual’s metabolic fate.

It is noticeable that, in young adulthood (prior to the HFD-challenge), although slight differences in body weight and triglyceride (TG) levels were observed between groups, all the biochemical parameters fell within the normal range and were not predictive of an individual’s sensitivity to DIO. We therefore attempted to identify, in this second experimental model, markers associated with significant changes in terms of susceptibility to obesity when animals are exposed to an obesogenic diet and that could be detected early (before the HFD-challenge).

### Identification of a set of genes whose expression correlates with predisposition to DIO in a nutritional programming model

RNA-seq analysis was conducted on pg-WAT samples from 5-month-old animals from groups A, B, and D that had not been exposed to HFD (Array Express E-MTAB-13879). The analysis workflow is illustrated in Figure 4A. First, EdgeR-based differential expression analysis was performed on the RNA-seq data, resulting in the identification of 2145 mRNA species that displayed significant differential expression for the group factor (A, B, D). This preliminary statistical analysis was later used to refine the list of genes. The same dataset was then subjected to PLS-DA to identify variables capable of distinguishing individuals between the three groups. As previously, to assess the reliability of the predictive model obtained and to determine if overfitting or excessive randomness were problematic, permutation tests were conducted (Figure S1C). The error rates obtained with the permuted data were higher than those obtained with the original dataset, confirming the validity of the predictive model. The contribution of each variable to the predictive model was assessed by calculating the VIP. In total, 1889 mRNA species exhibited a VIP ≥ 1.5. These corresponded to 1873 unique gene symbols, which were selected and considered as potential variables of interest in the analysis (Figure 4B).

**Figure 4:**
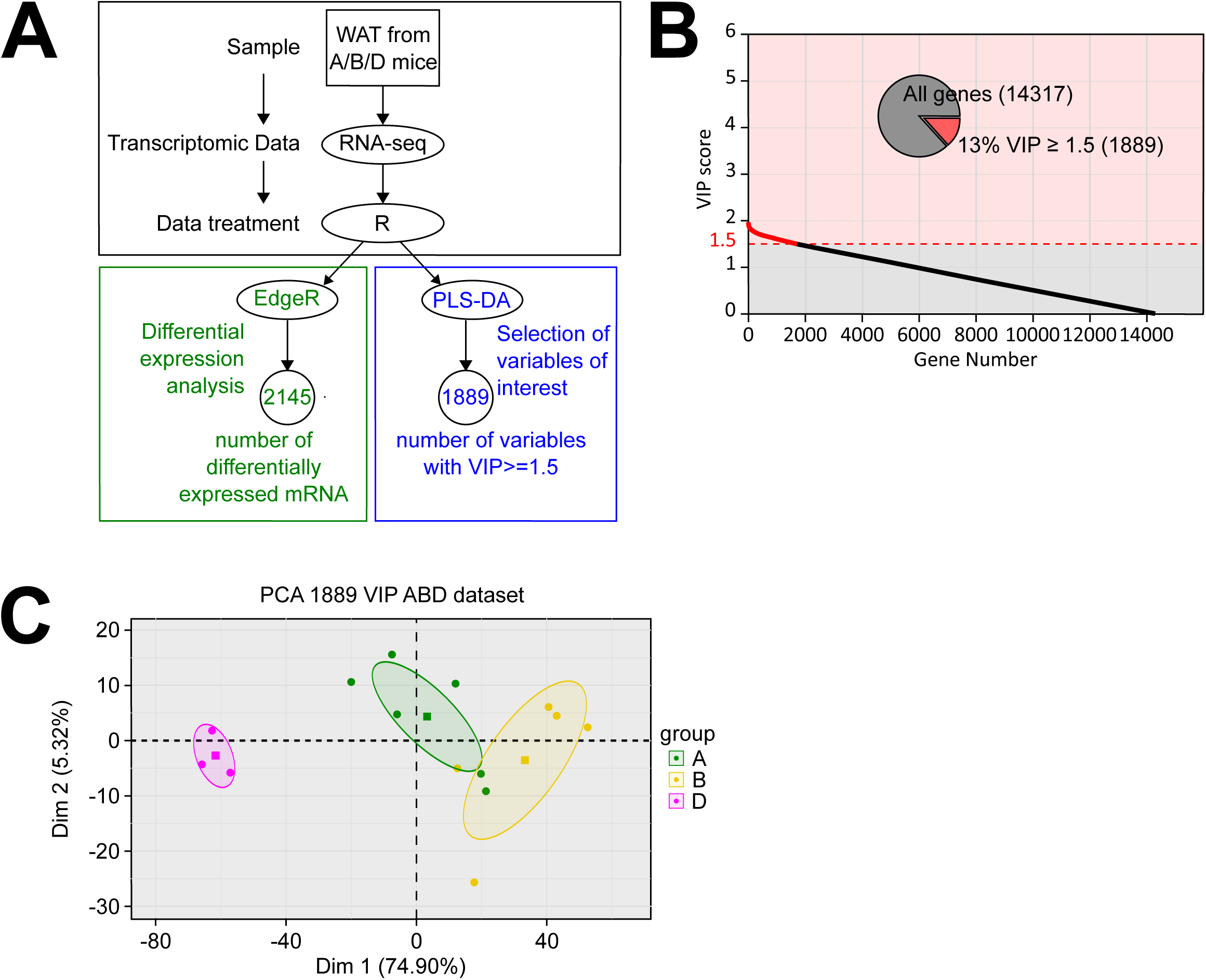
Identification of a set of genes whose expression correlates with predisposition to DIO in a nutritional programming model. **A: Workflow to select genes of interest.** RNA sequencing was performed on RNA extracted from pg-WAT harvested from 5-month-old mice from the three groups A (n= 6), B (n= 5), and D (n= 3). Raw data were treated with the R package EdgeR prior to differential expression analysis (green part of the workflow). The mixOmics R package was used for PLS-DA (blue part of the workflow) to identify important genes based on their VIP scores. **B: Variable of importance in projection (VIP) scores for each gene used in the PLS-DA.** Genes contributing meaningfully to the PLS-DA model with a VIP score >1.5 constitute 13% of the genes tested (inset pie-chart). The y-axis indicates the VIP scores for each variable indicated on the X-axis. Red dots indicate variables with the highest VIP scores (≥1.5) and thus contributing the most to class discrimination in the PLS-DA model. **C: Principal Component Analysis (PCA).** PCA was performed using the set of 1889 mRNA selected by PLS-DA. Data from groups A (green dots), B (yellow dots), and D (pink dots) are plotted along the first (X) and second (Y) principal components. Ellipses represent the 95% confidence interval and squares represent the barycenter of each group.

Using PCA (Figure 4C), we investigated how well these 1889 mRNA species classify mice in their original groups. The results revealed that these mRNA species were effective indicators of membership of either the B or D groups. Taken together, these results demonstrate that maternal nutritional stress can produce a metabolic imprint in offspring, making them more or less prone to DIO and related conditions. This imprint is associated with a specific gene expression pattern, which could potentially serve as a means to determine susceptibility to DIO.

### Common genes signature have discriminant potential regarding predisposition to DIO

To identify a robust set of predictive genes, we compared the gene signatures obtained from the PLS-DA applied to our two experimental models (Figure 5A). This comparison led to the identification of 201 common discriminant genes. However, upon further analysis, four of these genes (despite having a VIP ≥ 1.5 for the 3 analyses described above) were found to have an adjusted q-value (as calculated above using ANOVA) exceeding 0.05 across all three datasets (ABD, T0, T18). Consequently, these genes were excluded from the analysis. As a result, a final set of 197 genes was selected for further investigation (Table S1, column A).

**Figure 5:**
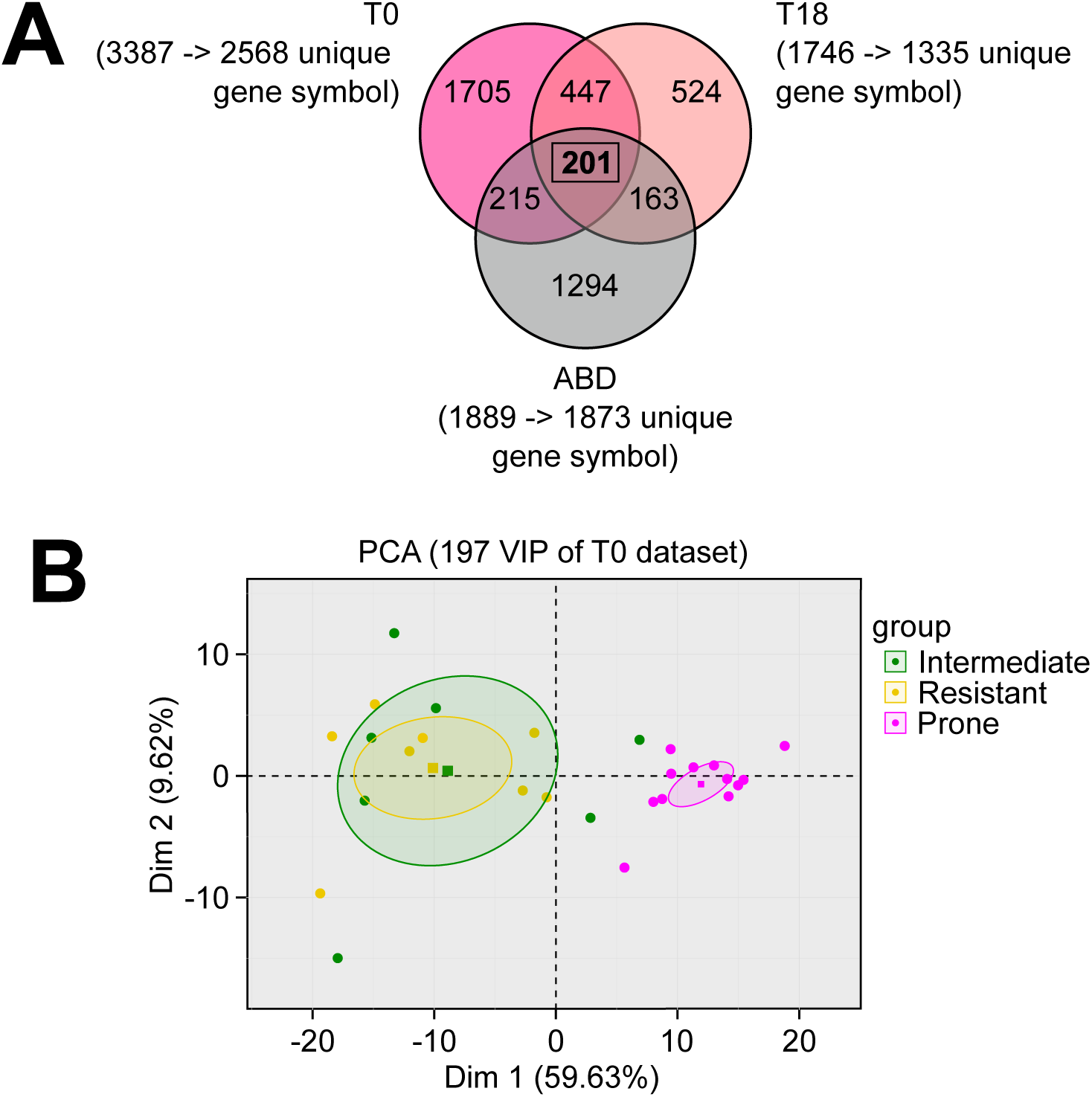
Identification of candidate genes and evaluation of their discriminant potential. **A: Identification strategy.** To identify the most robust predictive genes, we compared the three gene lists (containing 1889, 1746, and 3387 genes from the three datasets ABD, T0 and T18) obtained by PLS-DA and identified a list of 201 genes. Among these 201 common genes, 4 presented an adjusted q-value >= 0.05 for all three data sets (ABD, T0, T18) and were therefore eliminated, leaving 197 selected genes (Table S1, column A). **B: Principal Component Analysis (PCA)** was performed using the expression of the 197 selected genes from the first model at T0. Data from groups I (green dots), R (yellow dots), and P (pink dots) are plotted with respect to the first (X) and second (Y) principal components. Ellipses represent the 95% confidence interval and squares represent the barycenter of each group.

To assess the discriminatory potential of these 197 genes before the HFD-challenge, we performed PCA on T0 data from model 1. The results shown in Figure 5B demonstrate that, based on the expression patterns of these 197 genes in experimental model 1 at T0, it is indeed possible to distinguish obesity-prone animals from the animals in the other groups. This finding highlights the potential usefulness of these genes for the reliable identification of individuals with a predisposition to DIO.

### Candidate genes identified in mice also discriminate between lean and obese human individuals

As our long-term goal is to transpose the results obtained in mice to humans, it was important to know whether the human orthologs of the 197 candidate genes identified in our two mouse models above could also be considered as good candidates to be predictive of susceptibility to obesity in humans. To determine whether this was the case, we first had to address several important ancillary questions.

Our first concern was whether these markers were also detected in other WAT depots, such as subcutaneous WAT, as this type of deposit would be more accessible in humans. From an experimental point of view, it is very complicated to obtain subcutaneous WAT samples from mice, especially when working with lean mice.

Secondly, we questioned whether these markers could also apply to female animals. Even though this issue could have also been addressed in female mice, they are not an ideal experimental model for obesity susceptibility as they are mostly resistant to DIO ^18,19^.

Due to the difficulties inherent to mouse models, we considered it would be more relevant to address these two questions directly using human data. To do so, we opted to rely on publicly available datasets from human studies. Thus, to assess whether the 197 genes identified in mice might also be discriminative in humans, we combined our three mouse datasets, described above, with four independent external datasets derived from previous studies on humans ^20–23^. To closely match human data, the mouse data sets used were composed of groups A and D from the nutritional programing model and group I and P from the innate variability model at T0 and T18. Indeed, our goal is to identify individuals who are sensitive compared to the general population, rather than compared to a specific, resistant population.

Regarding the human datasets used, further details can be found in Table S2. Briefly, each dataset is composed of two groups (lean and obese) for which transcriptomic analysis (either through microarray or RNA-seq) was performed on WAT (either subcutaneous or omental), from both male and female individuals. For our analysis, we first analyzed all datasets to identify which of our 197 mouse genes are present in all human datasets. Following this alignment, 143 genes remained (Table S1, column B) for which expression data was available for a total of 141 individual samples (either mice or human) from the seven integrated datasets. It is important to note that the supplemental human datasets include numerous confounding co-factors such as ethnic origin, gender, age, adipose tissue deposit, that were not controlled in this analysis. Despite the potential effects of these confounding factors, all samples from the external studies were included, with the aim of producing robust predictive models that are not influenced by the potential effects of these factors. To deal with the heterogeneity due to the use of data from various sources, horizontal data integration was performed using the R mixOmics package MINT methods (P-integration). In the MINT toolbox, we used mint.plsda and mint.splsda to build two successive classification models for discriminative and predictive purposes. A first basic 5-component classification model, including the 143 common variables with a two-group outcome (lean vs obese) was produced, and its performance assessed. A first graphical output (Figure 6A) projected all the samples onto a unique space, with a good separation of samples based on their group, especially along the axis corresponding to component 1 (which retains 17% of the overall variance). A second visualization allowed the partial projection of samples, one study after another (Figure 6B). The results confirmed that the group discrimination is adequate for all seven studies, whether involving mice or humans, especially along the axis for component 1. This analysis also assigned weights (or loading) to the different variables depending on their relative contributions to the model (Figure 6C + Table S1, column C). Based on this loading, the candidate genes could be hierarchically organized. The 23 most predictive genes – those with the highest absolute values, at the top of the hierarchy – are presented in Figure 6C

**Figure 6:**
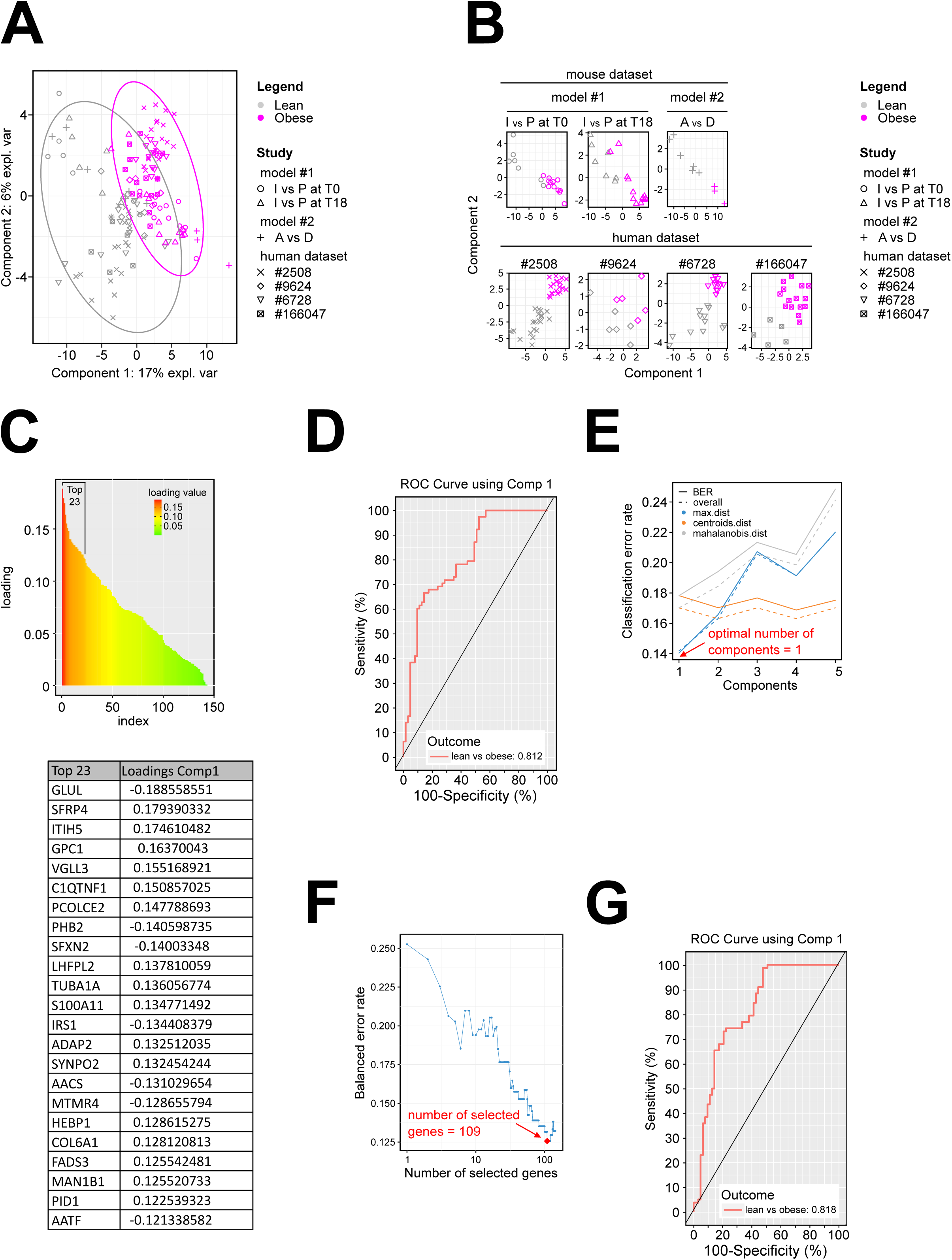
Candidate genes identified in mice discriminate between lean and obese humans Three mouse data sets were used: nutritional programing model (groups A and D), innate variability model at T0 (group I and P) and innate variability model at T18 (group I and P) together with four human datasets from publicly available resources (GEOD-2508: microarray data obtained from isolated adipocytes of subcutaneous adipose deposits from Pima Indians, male or female and lean or obese. Cohort MTAB-6728 contains a large number of lean and obese individuals without specifications on biological sex. Cohort GSE-166047 consists of lean and obese females. A cohort of children (E-GEOD-9624) for whom omental adipose tissue samples were analyzed, see Table S2 for details and references). (A-E) MINT.plsda (Multivariate INTegration plsda) including both mouse and human datasets. A: Sample plot from the MINT PLS-DA performed on the seven gene expression datasets. Samples are projected into the space corresponding to the first two components. Sample colors reflect their group (lean, obese), and symbols indicate the source study. B: Study-specific sample plots showing the projection of samples from each of the datasets in the same subspace spanned by the first two MINT components. C: Top, Distribution of genes according to their loading values. Bottom, top 23 of the genes according to their loading value (loading value > 0.12), selected arbitrarily, for illustration purpose, on the basis of the break observed in the distribution graph. D: ROC curve and AUC from the MINT PLS-DA for the seven gene expression datasets. In a ROC curve the true positive rate (i.e. the proportion of correctly predicted positive instances relative to all actual positive instances = Sensitivity) is plotted in function of the false positive rate (i.e. the proportion of incorrectly predicted positive instances relative to all actual negative instances = 100-Specificity). The numerical output indicated is the area under curve (AUC) measuring the overall performance of the model. It quantifies the ability of the model to discriminate between positive and negative classes. A higher AUC indicates better performance. AUC values greater than 0.8 are often considered very good and suggest a model with strong discriminative power. (E-G): MINT.splsda (Multivariate INTegration sparse plsda) including both mouse and human datasets. E: Choosing the number of components in mint.splsda using “perf()” with LOGOCV (Leave One Groupe Out Cross Validation) applied to the seven gene expression datasets. Classification error rates (overall and balanced) are represented on the y-axis with respect to the number of components on the x-axis for each prediction. The plot shows that the error rate reaches a minimum from one component with the BER and max distance. F: Tuning keepX in MINT.splsda performed on the seven gene expression datasets. The line represents the balanced error rate (y-axis) for component 1 across all tested keepX values (x-axis). Is called keepX the minimum number of gene necessary to keep a model with at least the same performance than the initial model. The diamond indicates the optimal keepX value which achieves the lowest classification error rate as determined with a one-sided t−test across the studies. G: ROC curve and AUC from the MINT.splsda performed on the seven gene expression datasets.

The performance of this first prediction model when seeking to distinguish between lean and obese individuals can be assessed in two ways: the first output is the ROC (Receiver Operating Characteristic) curve and its AUC (Area Under Curve) which give a first clue as to the overall quality of the predictive model ^24^ (Figure 6D). With an AUC at 0.812 when focused on component 1, the model has very good capacity to predict the two phenotypes. This was confirmed by the second output, which corresponds to the diverse low error rates calculated by the “perf” functions and its LOGOCV algorithm (Leave one Group Out Cross Validation). This algorithm sequentially isolates each of the 7 datasets and uses it as an external set to be predicted. It returns overall error rates of these predictions that assess the model’s predictive power. For our first predictive model, we looked mainly at the BER (Balanced Error Rate) on component 1, the main variability axis containing most of our data (Figure 6A). For this component, the BER value obtained – 0.178 – was satisfactory. Another parameter, namely the group error rate on component 1, gave a very satisfactory value of 0.10 for the prediction of the “obese” group, whereas a higher value of 0.254 was obtained for the “lean” group, indicating that this class is more difficult to assign (predict), probably due to the higher dispersion of its samples (Figure 6A).

Secondly, an optimized model was generated using the same data and the sparse mint.splsda method. This optimization process involved determining the minimal number of (1) components, and (2) genes that can be retained in the model without affecting prediction rates. Following this optimization, only one component was retained in the model (Figure 6E); it was associated with 109 of the initial 143 genes (Figure 6F). Applying these two parameters reduced the overall error rates (from 0.1783 to 0.1306). In comparison to the first model outputs, the gene hierarchy remained the same, but with higher absolute loading values (for the first 34 genes) (Table S1, column E). The quality assessments, from ROC curves (Figure 6G) or error rates, revealed a slightly better model with an AUC of 0.818, a BER at 0.131, and a group error rate of 0.16 for the lean group. The error rate for the obese group remained unchanged (0.1026).

Functional annotation of all 197 genes identified in mice is given Table S1 (columns K to AD). Based on this annotation, enrichment analysis was performed, using Metascape ^25^ for the 143 genes present in all mice and human datasets (Figure S4A) and for the optimized list of 109 genes (Figure S4B).

Our analysis revealed a hierarchically-ordered set of genes with expression patterns associated with (1) obesity in humans and (2) a predisposition to obesity in mice. Importantly the expression of this set of genes is found to be differentially expressed in mice that are predisposed to obesity, even before any clinical symptom arises. These genes are thus good candidates for exploring the possibility of developing a human diagnostic test to predict predisposition for obesity.

## DISCUSSION

We successfully identified a set of genes in WAT for which expression levels significantly correlated with susceptibility to obesity. To our knowledge, our study is the first to describe predictive markers of predisposition to obesity that are differentially expressed before the onset of obesity symptoms. Many studies, in both rodents and humans, have previously identified state markers of obesity ^33,34^. However, these state markers are not predictive but rather descriptive of an established condition. Notably, some are linked to maternal obesity ^35^, which is, in itself, predictive of offspring obesity. In our study, the mothers of predisposed offspring are not obese, and our predictive markers are therefore independent of the mother’s physiological state. Furthermore, many studies have identified genetic variants that predispose individuals to obesity ^36,37^, but it is important to note that genetic variants account for less than 5% of obesity cases (and they often lead to very severe, early-onset obesity). Most of obesity cases cannot be predicted using these genetic variants.

Two notable observations were made regarding the expression of the predictive set of genes identified: (1) When offspring are exposed to a maternal nutritional stress that increases theirs predisposition to obesity, the expression pattern for these genes is altered and remains perturbed throughout their lifetime. (2) In a cohort of mice, expression of these genes is systematically altered in animals that are naturally predisposed to obesity, even before the nutritional challenge (HFD feeding) leading to weight gain. It is likely that the causes of the observed natural variability are of a similar nature to those established in the perinatal nutritional stress model, but that they occur stochastically, depending for example on the size of the litter, the position of the fetus in the uterine horn, maternal care, etc…) rather than being induced by maternal nutritional stress. This appears to suggest that natural imprinting, influenced by various factors, contributes to the variability in phenotypes observed even among individuals within the same experimental group.

An important question is how expression of the set of genes identified is regulated. Our observations suggest that the mechanisms at play could involve epigenetic mechanisms that operate separately from genetic factors. Indeed, due to the use of an inbred mouse strain, there should be no major genetic differences between animals in our cohort. The consequences of this programming may remain latent and only manifest under specific conditions, such as when exposed to a HFD or later in the aging process. At this stage of our studies, we do not know whether the epigenetic modifications directly affect the expression of the identified genes or if they converge on one or more common effectors. Such epigenetic modifications are likely to be complex to detect. Methylation at CpG sites and post-translational modifications of histones may play a pivotal role in this context. Among these epigenetic modifications, we investigated the methylome of these mice; however, it was very difficult to obtain precise measurements, and the data were less accurate than anticipated, making it impossible to draw robust conclusions. It is also noteworthy that among the 197 predictive markers identified in mice, two are known epigenetic factors: HDAC9 (histone deacetylase) and PHF8 (a histone demethylase that specifically demethylates H3K9me2/1). Their roles in regulating the expression of the other markers warrant further investigation.

Regardless of its fundamental nature, the imprint contributing to susceptibility to DIO appears to be remarkably stable. Indeed, it is noticeable that markers showing a correlation with sensitivity to DIO were consistently observed at different ages and stages of obesity progression. Nevertheless, while this study did not explore the potential reversibility of these stable markers, it is conceivable that long-term interventions, such as healthy eating or physical activity, could potentially mitigate or ameliorate the predisposition to obesity due to the early developmental conditions to which the individuals were exposed.

Assuming that predisposition to obesity is determined epigenetically, it would be interesting to determine when this modification takes place. Our findings suggest that, based on biological parameters (data not shown), consumption of a LPD during lactation (group E) or during gestation + lactation (group B) results in a similar phenotype. This suggests that the lactation period plays a significant role in shaping outcome for pups. It should be noted that, in mice, this period corresponds to the period of adipose tissue development. In rodents, WAT is absent at birth and develops in the first 2 weeks *ex utero*. In contrast, in humans, WAT deposits develop mainly during the last trimester of gestation. As a consequence, when comparing humans and rodents in terms of adipose tissue development, the lactational period in rodents may be more equivalent to the third trimester of human pregnancy. Later in life, WAT expands with age in most mammalian species, with relatively similar functions and locations of WAT ^26^.

Changes in WAT directly influence body weight by regulating energy storage and release, altering metabolic function, and secreting hormones that control appetite and insulin sensitivity. Excess WAT accumulation leads to obesity, while efficient fat breakdown promotes weight loss. The balance between fat storage and utilization, along with the endocrine activity of WAT, makes it a central player in body weight regulation. WAT biology and how it affects obesity development also present major sex differences in terms of composition and location. In mice, when consuming a HFD, females tend to exhibit a higher resistance to obesity than males^18,19^ (interestingly, this protection is reversed after ovariectomy ^27^). For this reason, we decided to explore sex differences in humans using publicly available data. The human data integrations were designed to challenge our selection of 197 mouse genes in a human context, including both sexes and different WAT deposits. The two main goals were to assess the discriminant potential of our selected genes and to develop predictive classification models with a larger scope. One of the major advantages of the MINT methods used is that they can cope with heterogeneity across several datasets. They assess and control for specific variations within each study to better reveal a common signature. As a result, confounding co-factors present in some datasets may not prevent the discovery of a set of discriminant variables applicable across all studies. With this set of tools, we first ranked the discriminant genes that separated the whole samples by class in each study. Then, a classification model based on these genes was established and its performance evaluated for the prediction of the status of new samples. This first model exhibited promising characteristics, identifying a discriminant set of genes applicable to both mouse and human samples. Establishing a hierarchy among variables using loadings calculated across various studies provides a robust foundation for the selection of the best markers. The evaluation of the model’s predictive potential, particularly focusing on the BER along axis 1, suggests a reasonably low error rate. Despite using a relatively large number of genes (n = 143), the low error rates across all the axes evaluated indicate that the selected variables do not introduce significant noise into the analysis. Consequently, the initial gene selection appears to be applicable and transferable to human contexts, making the model adaptable to assess predisposition to obesity in humans.

The second MINT.splsda model (Figure 6F-G) was optimized for its capacity to identify the group of individuals using a minimal number of components and a minimal number of genes. Remarkably, 109 out of the initial 143 genes were required to obtain accurate predictions. This suggests that the expression patterns of the majority of the initial gene set contribute valuable and non-redundant information, emphasizing the importance of considering each variable when seeking precise predictions. The conservation of marker hierarchy from the first model, despite changes in absolute values, underlines the stability of the selected marker set.

It is noticeable that the three highest-ranking genes – Glul, Sfrp4, and Itih5 – have been previously linked to metabolic diseases. Itih5 (inter-α trypsin inhibitor H5) belongs to the inter-α trypsin inhibitor heavy chain (ITIH) gene family, encoding secreted heavy chain peptides ^28^. This gene is expressed at high levels in adipose tissue, and its expression is higher in obese subjects than in lean subjects. Its expression reduces during diet-induced weight loss ^29^. Sfrp4 (Secreted frizzled-related protein 4) belongs to the SFRP family of secreted glycoproteins. These soluble modulators are believed to alter Wnt signaling ^30^. SFRP4 is the largest member of the SFRP family and has been implicated in glucose and lipid metabolism following interaction with Wnt ligands. Its level is increased in obesity ^31^. Glul encodes glutamine synthetase, which uses glutamate as a substrate and is the only known glutamine-synthesizing enzyme. Petrus et al. ^32^ showed that glutamine metabolism is perturbed in WAT from obese individuals and that Glul was one of the most prominently dysregulated genes in the glutamine pathway in these samples.

Furthermore, there are several avenues for refinement, facilitation, and enhancement of the predictive capabilities of this set of genes in human. Firstly, it would be necessary to obtain access to human cohorts with longitudinal follow-up, with access to adipose tissue biopsies collected over multiple years and including data tracking changes in patient weight. Secondly, the effective markers identified could be further investigated for their presence as circulating transcripts or proteins, making them more accessible for predictive and diagnostic purposes. Nevertheless, since these markers can also be effectively monitored in subcutaneous WAT, a biopsy could be envisaged for diagnostic purposes. Thirdly, the predictive potential of these markers regarding susceptibility to the development of insulin resistance and type 2 diabetes remains to be determined. This could be possible in humans, where clinical parameters relative to insulin sensitivity are easier to measure than in mice. The gene signature identified and its predictive capabilitie offers a promising means to address the complex biological phenomena that is obesity. Ultimately, the data obtained take us a step closer to personalized medicine. By detecting at-risk individuals in the early stages, it may be possible to offer advice on nutrition and physical activity to mitigate the negative metabolic impact associated with their intrinsic predisposition.

## AUTHOR CONTRIBUTIONS

Conceptualization: CJ, PF; Data curation: GC; Formal Analysis: CJ, GC, MB, JT; Funding acquisition: PF, CJ; Investigation: CJ, LP, YM; Methodology: CJ, LP, PF; Project administration: CJ, PF; Visualization: CJ, GC; Writing – original draft: CJ, PF, GC; Writing – review & editing: CJ, PF, GC, AB, ACM, JA, CV.

## ACKNOWLEDGMENTS

The authors thank UNH’s animal housing facility (IEN, Julien Hermet), Yoann Delorme, Mehdi Djelloul-Mazouz for help with animal work and Anne Terrisse for her involvement in animals welfare, sequencing facilities IGENSEQ (Yannick Marie, ICM Institute, Paris, France) and GeT-TRiX (Yannick Lippi, GénoToul, Génopole Toulouse Midi-Pyrénées). The authors also thank Dr. Lydie Combaret, Daniel Taillandier, Cécile Polge, Etienne Lefay, Marie Lefebvre (UNH, INRAe Theix), Patrick Even (INRAe, Paris) and Ez-Zoubir Amri (université Côte d’Azur, Nice) for constructive discussions and Mélanie Vachette-Dit-martin for participating to experimentation. We also thank Dr Maighread Gallagher for proofreading the English of our manuscript. This work was supported by Agence Nationale de la Recherche (Epidiabesity and Epimetab grants).

## DECLARATION OF GENERATIVE AI AND AI-ASSISTED TECHNOLOGIES IN THE WRITING PROCESS

During the preparation of this work the authors used Chatgpt in order only to improve readability and language of the manuscript. After using this tool, the authors reviewed and edited the content as needed and take full responsibility for the content of the publication.

## STAR METHODS

### RESOURCE AVAILABILITY

#### Lead contact

Further information and requests for resources should be directed to and will be fulfilled by the lead contact, Céline Jousse

#### Materials availability

This study did not generate new unique materials.

#### Data and code availability

- Data:
  - Transcriptomic data reported in this paper have been deposited at ArrayExpress. The ArrayExpress accession numbers are listed in the key resources table.
  - This paper analyzes existing, publicly available data. These accession numbers for the datasets are listed in the key resources table
- Code:
  - This paper does not report original code
- Any additional information required to reanalyze the data reported in this paper is available from the lead contact upon request.

## EXPERIMENTAL MODEL

All experiments were conducted with the approval of the regional ethics committee (agreement no. D6334515) following the European Directive 2010/63/EU on the protection of vertebrate animals used for experimental and scientific purposes.

All BALB/c mice (strain BALB/cAnNRj) were purchased from Janvier-Labs (Le Genest-Saint-Isle, France) at 5-weeks of age. Upon arrival, they were housed in a controlled room (22 ± 2◦C, 60 ± 5% humidity, 12 h light/dark cycle, and light period starting at 8:00am), fed *ad libitum* a standard rodent diet (A03 from Safe, Augy, France), and given free access to water, until they reach the age required for experimental protocols described below.

### Model 1

Twenty-seven 4-month-old BALB/c male mice were used. At T0, perigonadal WAT (pgWAT) biopsies from each mouse were collected. One week after surgery, mice were fed an experimental High Fat Diet (HFD, Open Source Diet, Ref D12451) for 18 weeks after which they were sacrificed (T18) for removal of perigonadal WAT. The experimental plan is outlined in Figure 1A. Body weight and body composition were monitored at various stages. The initial number of animals (n=27) was determined based on previous experiments in order to be able to generate 3 groups of n >6 mice based on body weight changes following HFD challenge (either Resistant, Intermediate or Prone to DIO)

### Model 2

Two-month-old virgin BALB/c female mice fed a chow diet (A03; Safe, Augy, France) were mated with 2-month-old BALB/c male, isolated at vaginal plug detection and randomly transferred on Experimental Diet (CD: Control Diet, LPD: Low Protein Diet, HFD: High Fat Diet) as indicated figure 3A (n= 5, 8, 11, 3 and 10 dams respectively for each of the experimental groups A, B, C, D and E). At parturition, randomization was carried out as follows: dams and litters were transferred on experimental diet as indicated figure 3A. Litters of different sizes were obtained in each group of pregnant female. Since the litter size is important in the offspring life trajectory, our strategy was to erase all possible confounding factors coming from litter size and the allotting protocol was as follows: Mice from litter of 3 pups or less or from litter of more than 10 pups were excluded to minimize the variability of each parameter known to be impacted by litter size and we thus considered only litters that have a total number of pups comprised between 4 and 10 to avoid extreme litter size; After weaning at 4 weeks of age, male offspring from each group were housed individually with free access to Control Diet (CD). To obviate any litter effects, animals used for each experiment were randomly chosen in different litters, and only a limited number of animals (n = 1 to 2) was used from each litter. The initial number of animal used was determined based on previous experiment in order to generate male offspring number > 8 per group.

CD and LPD (UPAE, Jouy-en-Josas, France) contain respectively 22% and 10% protein and composition are given ^11^. HFD (High Fat Diet) contains 45% kCal from fat (Open Source Diet Ref D12451).

Body weight and body composition were monitored at various ages throughout life for all F1 male mouse groups. Part of F1 male mice of each group were sacrificed at 6 mo for further analyses and part of F1 male mice from groups A, B and D were submitted to a HFD-challenge from the age of 5 to 12-month.

## METHOD DETAILS

### Body Composition

Individual mice were placed into restrain tube and inserted into the mouse EchoMRI-100 instrument (Echo Medical Systems LLC, Houston TX, USA) to determine both fat and lean mass (g). Total body weight was measured using a standard top-loading laboratory balance. Adiposity is expressed as a percentage of fat relative to total body weight.

### Metabolic parameters

Total cholesterol, HDL-cholesterol, triglycerides, and glucose concentration were measured in plasma of over-night starved adult male mice, using commercial enzymatic assays. For oral glucose tolerance test (OGTT) mice were given an oral glucose bolus (2 g/kg) after a 7-hour fast. Blood was collected from the tail vein at different times. Glycemia was measured using the OneTouch glucometer (Lifescan Inc., Milpitas, CA).

### Tissue collection

For perigonadal WAT biopsy, mice were placed on an heating-pad under isoflurane anesthesia using a nose cone. At the end of experiments, mice were euthanized by cervical dislocation under isoflurane anesthesia. In both cases, perigonadal WAT was collected prior to immediate freezing in liquid nitrogen and stored at −80 ◦C until analyses. Chirurgy and euthanasy were performed by alterning one mouse of each group in order to minimize potential confounders.

### Transcriptomic analysis

#### RNA-Seq gene expression studies

Prepared RNAs sample were sent to the sequencing facility (IGENSEQ) at the ICM Institute (Paris, France), and their quality and concentration were checked using bioanalyzer. Libraries were prepared using the “TruSeq Stranded mRNA” kit (ILLUMINA®), allowing the construction of a library of polyadenylated RNAs (mRNA and polyadenylated non-coding RNAs such as lncRNA) from total RNAs. The libraries were prepared following the manufacturer’s recommendations and subsequently sequenced using the Nextseq 500 (ILLUMINA®) to obtain 2 * 40 million 75- base pair fragments per sample.

#### Micro-array Gene Expression studies

Gene expression profiles were performed at the GeT-TRiX facility (GénoToul, Génopole Toulouse Midi-Pyrénées) using Agilent Sureprint G3 Mouse GE v2 microarrays (8×60K, design 074809) following the manufacturer’s instructions. For each sample, Cyanine-3 (Cy3) labeled cRNA was prepared from 50 ng of total RNA using the One-Color Quick Amp Labeling kit (Agilent Technologies) according to the manufacturer’s instructions, followed by Agencourt RNAClean XP (Agencourt Bioscience Corporation, Beverly, Massachusetts). Dye incorporation and cRNA yield were checked using DropsenseTM 96 UV/VIS droplet reader (Trinean, Belgium). 600 ng of Cy3-labelled cRNA were hybridized on the microarray slides following the manufacturer’s instructions. Immediately after washing, the slides were scanned on Agilent G2505C Microarray Scanner using Agilent Scan Control A.8.5.1 software and fluorescence signal extracted using Agilent Feature Extraction software v10.10.1.1 with default parameters.

## QUANTIFICATION AND STATISTICAL ANALYSIS

Cohen’s effect size was calculated as followed: 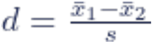 with 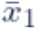 = mean of Group 1, 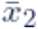 = mean of Group 2, *s* = standard deviation

### Editing and statistical analysis of mouse datasets

Data editing and statistical analysis have been performed thanks to R sofware in its 4.2.3 version, in Rstudio (2022.07.2) and GraphPad.

For mouse microarray data, differential analysis between the three groups (R, I, P) was performed with the limma package (v3.54.2) through the lmFit() and eBayes() functions ^38^. A selection threshold was applied at 0.05 on the Benjamini-Hochberg adjusted p-values.

For mouse RNA-Seq data, the standard edgeR (v3.40.2) ^39^ procedure was applied to clean and transform the data. Genes with a cpm transformed counts mean below 10 were first rejected from analysis as low expressed. Before differential analysis a TMM (trimmed mean of M values) normalisation procedure was applied on filtered raw counts to compensate technical bias at the samples level. EdgeR quasi-likelihood F-test was then used to select significant genes with a Benjamini-Hochberg adjusted p-values below a 0.05 threshold. Further analysis were applied on log(cpm) transformed and thus linearised counts data.

### Principal Component Analysis

The discriminant capacity of the genes lists obtained were tested by PCA using the R FactomineR package on various mouse and human datasets.

### Hierachical Clusturing Analysis

The hclust function was used to perform hierarchical clustering (based on a Ward.D2 aggregation algorithm) in order to build a classification based on physical parameters measured at T18.

### PLS-DA

Partial Least Square discriminant analyses was used to identify the important variables (i.e., mRNA) for group classification (R package mixOmics) ^40^. PLS-DA is a PLS approach in which only one dependent variable represents the class or group membership. The variable influence on projection (VIP) indicates the influence of each variable on the discrimination between the different groups. For each model, the most important transcripts in the classification were ranked according to decreasing VIP. Genes with a VIP score >1.5 were considered as contributing meaningfully to the PLS-DA model. For each PLS-DA model built, a permutation test was realized to evaluate whether the observed discrimination between groups is statistically significant or could have occurred by chance. The permutation test for PLS-DA involves randomly permuting the group membership of the observations and recalculating the PLS-DA model for each permutation. The performance of each of the model obtained are assessed by their classification error rate and compared to the error rate of the true model. This procedure is repeated 500 times.

### P-integration : Mixomics MINT-PLSDA

In order to combine our own mouse data with external datasets and to better evaluate the discriminant potential of our markers, an horizontal integration approach (P-integration) was implemented thanks to the R mixOmics package (v6.22.0) and its MINT functions (http://mixomics.org/mixmint) ^41,42^. This MINT method is dedicated to the integration of multiple datasets, from multiple sources, measured on the same set of genes, thus quantified in different context (studies). Here supervised pls-da tools have been used for modelisation. Basically, 5 components models have been created. The samples have been projected in factorial plans to assess the groups separation globally and at the study level. The genes contributions to the 2 first factorial axis have also been calculated. ROC curves and their area were used as first indicator of model’s quality. Finally using one by one human datasets for external validation, we were able to calculate new ROC curves, this time associated with error rate and balanced error rate values, adding an increased confidence in our integrative classification models.

Seven datasets have been combined : our 3 mouse datasets with 4 human ones. All have been aligned at the features (variables) level, from the 197 genes list. This alignment kept 143 variables quantified for 141 samples corresponding to the whole cases (individuals) from the 7 integrated datasets. These samples where annotated as either « lean » or « obese » to characterise the two physiological states under study. This gives a global vector used as a target to define the classes to discriminate. A last vector defining the studies blocks was added to split the fusion dataframe into the seven experimental context and fullfill the functions options. The supplemental human datasets include potential confusing co-factors such as specie, gender, age, tissue, that were not controlled in this analysis. Despite these potential effects, all the external studies samples were included, with the aim of gaining robust models overwhelming potential side effects.

In MINT toolbox we used the supervised framework with its mint.plsda et mint.splsda to build two succeccive classification models. The pls algorithm is especially useful in the case of a limited number of samples in comparison to the number of quantified features. A first basic model including all the common variable, without any kind of further selection, based on 5 components was produced and its performance assessed. Secondly, an optimised model was released through the sparse method, that reduced model’s dimensions to a single component and the diversity of features to 109 genes from the 143 initial ones. These two models help in discriminative and predictive purposes.

### Functional enrichment analysis

Online Metascape (http://metascape.org, v3.5) ^25^ allowed the functional annotation of our gene sets. Custom analysis used a specific background derivated from commun transcriptome between all the human and mice datasets.

## SUPPLEMENTAL INFORMATION

Document “suppl figures”: Figure S1-S4. Figure S1: Additional permutation test related to Figures 1,2,4. Figure S2: Plasma metabolites parameters of 5-mo-old male mice from group A, B, and D, related to figure 3. Figure S3: Body weight and fat mass change following a 7-months HFD-challenge on groups A, B and D, related to figure 3D. Figure S4: Gene Ontology (GO) enrichment analysis related to figure 6

Table S1: Excel file containing additional data that are too large to fit in a PDF. Annotation and GO-term summary for the lists of genes identified along the analyses.

Table S2: Excel file containing additional data that are too large to fit in a PDF. Description and references for the human datasets used

## ABBREVIATIONS

AUC: Area Under Curve
CD: Control Diet
DIO: Diet-induced-obesity
DOHaD: Developmental Origins of Health and Disease
HCA: Hierarchical Clustering Analysis
HFD: High-Fat Diet
LPD: Low Protein Diet
LOGOCV: Leave One Group Out Cross Validation
OGTT: oral glucose tolerance test
PCA: Principal Component Analysis
pg-WAT: perigonadal White Adipose Tissue
PLS-DA: partial least squares discriminant analysis
ROC: Receiver Operating Characteristic
T2D: Type 2 Diabetes
TG: triglyceride
VIP: Variable Importance in Projection

## REFERENCES

1. Billings, L.K., and Florez, J.C. (2010). The genetics of type 2 diabetes: what have we learned from GWAS? Annals of the New York Academy of Sciences 1212, 59–77. 10.1111/j.1749-6632.2010.05838.x.

2. Huang, L.-T. (2020). Maternal and Early-Life Nutrition and Health. IJERPH 17, 7982. 10.3390/ijerph17217982.

3. Barker, D.J., and Osmond, C. (1986). Infant mortality, childhood nutrition, and ischaemic heart disease in England and Wales. Lancet 1, 1077–1081. 10.1016/s0140-6736(86)91340-1.

4. Barker, D.J., Osmond, C., Golding, J., Kuh, D., and Wadsworth, M.E. (1989). Growth in utero, blood pressure in childhood and adult life, and mortality from cardiovascular disease. BMJ 298, 564–567. 10.1136/bmj.298.6673.564.

5. Barker, D.J.P. (2007). The origins of the developmental origins theory. Journal of Internal Medicine 261, 412–417. 10.1111/j.1365-2796.2007.01809.x.

6. Chavatte-Palmer, P., Tarrade, A., and Rousseau-Ralliard, D. (2016). Diet before and during Pregnancy and Offspring Health: The Importance of Animal Models and What Can Be Learned from Them. IJERPH 13, 586. 10.3390/ijerph13060586.

7. Barker, D.J.P. (1995). Intrauterine programming of adult disease. Molecular Medicine Today 1, 418–423. 10.1016/S1357-4310(95)90793-9.

8. Barker, D.J.P. (2007). Obesity and early life. Obesity Reviews 8, 45–49. 10.1111/j.1467-789X.2007.00317.x.

9. Burcelin, R., Crivelli, V., Dacosta, A., Roy-Tirelli, A., and Thorens, B. (2002). Heterogeneous metabolic adaptation of C57BL/6J mice to high-fat diet. American Journal of Physiology-Endocrinology and Metabolism 282, E834–E842. 10.1152/ajpendo.00332.2001.

10. Levin, B.E., and Dunn-Meynell, A.A. (2000). Defense of body weight against chronic caloric restriction in obesity-prone and -resistant rats. American Journal of Physiology-Regulatory, Integrative and Comparative Physiology 278, R231–R237. 10.1152/ajpregu.2000.278.1.R231.

11. Jousse, C., Parry, L., Lambert-Langlais, S., Maurin, A., Averous, J., Bruhat, A., Carraro, V., Tost, J., Letteron, P., Chen, P., et al. (2011). Perinatal undernutrition affects the methylation and expression of the leptin gene in adults: implication for the understanding of metabolic syndrome. FASEB j. 25, 3271–3278. 10.1096/fj.11-181792.

12. Jousse, C., Muranishi, Y., Parry, L., Montaurier, C., Even, P., Launay, J.-M., Carraro, V., Maurin, A.-C., Averous, J., Chaveroux, C., et al. (2014). Perinatal Protein Malnutrition Affects Mitochondrial Function in Adult and Results in a Resistance to High Fat Diet-Induced Obesity. PLoS ONE 9, e104896. 10.1371/journal.pone.0104896.

13. Dorlinga, M.W., Pawlakb, D.B., Ozanne, S.E., and Petry, C., J. Diabetes in Old Male Offspring of Rat Dams Fed a Reduced Protein Diet. INTERNATIONAL JOURNAL OF EXPERIMENTAL DIABETES RESEARCH.

14. Ikenasio-Thorpe, B.A., Breier, B.H., Vickers, M.H., and Fraser, M. (2007). Prenatal influences on susceptibility to diet-induced obesity are mediated by altered neuroendocrine gene expression. Journal of Endocrinology 193, 31–37. 10.1677/joe.1.07017.

15. Jones, A.P., Simson, E.L., and Friedman, M.I. (1984). Gestational Undernutrition and the Development of Obesity in Rats. The Journal of Nutrition 114, 1484–1492. 10.1093/jn/114.8.1484.

16. Alejandro, E.U., Jo, S., Akhaphong, B., Llacer, P.R., Gianchandani, M., Gregg, B., Parlee, S.D., MacDougald, O.A., and Bernal-Mizrachi, E. (2020). Maternal low-protein diet on the last week of pregnancy contributes to insulin resistance and β-cell dysfunction in the mouse offspring. American Journal of Physiology-Regulatory, Integrative and Comparative Physiology 319, R485–R496. 10.1152/ajpregu.00284.2019.

17. Desai, M., Byrne, C.D., Meeran, K., Martenz, N.D., Bloom, S.R., and Hales, C.N. (1997). Regulation of hepatic enzymes and insulin levels in offspring of rat dams fed a reduced-protein diet. American Journal of Physiology-Gastrointestinal and Liver Physiology 273, G899–G904. 10.1152/ajpgi.1997.273.4.G899.

18. Nishikawa, S., Yasoshima, A., Doi, K., Nakayama, H., and Uetsuka, K. (2007). Involvement of Sex, Strain and Age Factors in High Fat Diet-Induced Obesity in C57BL/6J and BALB/cA Mice. Exp. Anim. 56, 263–272. 10.1538/expanim.56.263.

19. Hwang, L.-L., Wang, C.-H., Li, T.-L., Chang, S.-D., Lin, L.-C., Chen, C.-P., Chen, C.-T., Liang, K.-C., Ho, I.-K., Yang, W.-S., et al. (2010). Sex Differences in High-fat Diet-induced Obesity, Metabolic Alterations and Learning, and Synaptic Plasticity Deficits in Mice. Obesity 18, 463–469. 10.1038/oby.2009.273.

20. Lee, Y.H., Nair, S., Rousseau, E., Tataranni, P.A., Bogardus, C., Permana, P.A., Allison, D.B., and Page, G.P. (2006). Microarray profiling of isolated abdominal subcutaneous adipocytes from obese vs non-obese Pima Indians: increased expression of inflammation-related genes.

21. Aguilera, C., Gomez-Llorente, C., Tofe, I., Gil-Campos, M., Cañete, R., and Gil, Á. (2015). Genome-Wide Expression in Visceral Adipose Tissue from Obese Prepubertal Children. IJMS 16, 7723–7737. 10.3390/ijms16047723.

22. Bjune, J.-I., Haugen, C., Gudbrandsen, O., Nordbø, O.P., Nielsen, H.J., Våge, V., Njølstad, P.R., Sagen, J.V., Dankel, S.N., and Mellgren, G. (2019). IRX5 regulates adipocyte amyloid precursor protein and mitochondrial respiration in obesity. Int J Obes 43, 2151–2162. 10.1038/s41366-018-0275-y.

23. Rey, F., Messa, L., Pandini, C., Launi, R., Barzaghini, B., Micheletto, G., Raimondi, M.T., Bertoli, S., Cereda, C., Zuccotti, G.V., et al. (2021). Transcriptome Analysis of Subcutaneous Adipose Tissue from Severely Obese Patients Highlights Deregulation Profiles in Coding and Non-Coding Oncogenes. IJMS 22, 1989. 10.3390/ijms22041989.

24. Au, E.H., Francis, A., Bernier-Jean, A., and Teixeira-Pinto, A. (2020). Prediction modeling—part 1: regression modeling. Kidney International 97, 877–884. 10.1016/j.kint.2020.02.007.

25. Zhou, Y., Zhou, B., Pache, L., Chang, M., Khodabakhshi, A.H., Tanaseichuk, O., Benner, C., and Chanda, S.K. (2019). Metascape provides a biologist-oriented resource for the analysis of systems-level datasets. Nat Commun 10, 1523. 10.1038/s41467-019-09234-6.

26. Rodgers, A., and Sferruzzi-Perri, A.N. (2021). Developmental programming of offspring adipose tissue biology and obesity risk. Int J Obes 45, 1170–1192. 10.1038/s41366-021-00790-w.

27. Grove, K.L., Fried, S.K., Greenberg, A.S., Xiao, X.Q., and Clegg, D.J. (2010). A microarray analysis of sexual dimorphism of adipose tissues in high-fat-diet-induced obese mice. Int J Obes 34, 989–1000. 10.1038/ijo.2010.12.

28. Zhuo, L., and Kimata, K. (2008). Structure and Function of Inter-α-Trypsin Inhibitor Heavy Chains. Connective Tissue Research 49, 311–320. 10.1080/03008200802325458.

29. Anveden, Å., Sjöholm, K., Jacobson, P., Palsdottir, V., Walley, A.J., Froguel, P., Al-Daghri, N., McTernan, P.G., Mejhert, N., Arner, P., et al. (2012). ITIH-5 Expression in Human Adipose Tissue Is Increased in Obesity. Obesity 20, 708–714. 10.1038/oby.2011.268.

30. Shi, Y., He, B., You, L., and Jablons, D.M. (2007). Roles of secreted frizzled-related proteins in cancer. Acta Pharmacologica Sinica 28, 1499–1504. 10.1111/j.1745-7254.2007.00692.x.

31. Guan, H., Zhang, Y., Gao, S., Bai, L., Zhao, S., Cheng, X.W., Fan, J., and Liu, E. (2018). Differential Patterns of Secreted Frizzled-Related Protein 4 (SFRP4) in Adipocyte Differentiation: Adipose Depot Specificity. Cell Physiol Biochem 46, 2149–2164. 10.1159/000489545.

32. Petrus, P., Lecoutre, S., Dollet, L., Wiel, C., Sulen, A., Gao, H., Tavira, B., Laurencikiene, J., Rooyackers, O., Checa, A., et al. (2020). Glutamine Links Obesity to Inflammation in Human White Adipose Tissue. Cell Metabolism 31, 375–390.e11. 10.1016/j.cmet.2019.11.019.

33. Hua, Y., Xie, D., Zhang, Y., Wang, M., Wen, W., and Sun, J. (2023). Identification and analysis of key genes in adipose tissue for human obesity based on bioinformatics. Gene 888, 147755. 10.1016/j.gene.2023.147755.

34. Liu, X., Sun, H., Zheng, L., Zhang, J., Su, H., Li, B., Wu, Q., Liu, Y., Xu, Y., Song, X., et al. (2024). Adipose-derived miRNAs as potential biomarkers for predicting adulthood obesity and its complications: A systematic review and bioinformatic analysis. Obes Rev 25, e13748. 10.1111/obr.13748.

35. Sharp, G.C., Lawlor, D.A., Richmond, R.C., Fraser, A., Simpkin, A., Suderman, M., Shihab, H.A., Lyttleton, O., McArdle, W., Ring, S.M., et al. (2015). Maternal pre-pregnancy BMI and gestational weight gain, offspring DNA methylation and later offspring adiposity: findings from the Avon Longitudinal Study of Parents and Children. Int J Epidemiol 44, 1288–1304. 10.1093/ije/dyv042.

36. Andersen, M.K., and Sandholt, C.H. (2015). Recent Progress in the Understanding of Obesity: Contributions of Genome-Wide Association Studies. Curr Obes Rep 4, 401–410. 10.1007/s13679-015-0173-8.

37. Loos, R.J.F. (2012). Genetic determinants of common obesity and their value in prediction. Best Practice & Research Clinical Endocrinology & Metabolism 26, 211–226. 10.1016/j.beem.2011.11.003.

38. Ritchie, M.E., Phipson, B., Wu, D., Hu, Y., Law, C.W., Shi, W., and Smyth, G.K. (2015). limma powers differential expression analyses for RNA-sequencing and microarray studies. Nucleic Acids Research 43, e47–e47. 10.1093/nar/gkv007.

39. Robinson, M.D., McCarthy, D.J., and Smyth, G.K. (2010). edgeR : a Bioconductor package for differential expression analysis of digital gene expression data. Bioinformatics 26, 139–140. 10.1093/bioinformatics/btp616.

40. Lê Cao, K.-A., and Welham, Z.M. (2021). Multivariate Data Integration Using R: Methods and Applications with the mixOmics Package 1st ed. (Chapman and Hall/CRC) 10.1201/9781003026860.

41. Rohart, F., Eslami, A., Matigian, N., Bougeard, S., and Lê Cao, K.-A. (2017). MINT: a multivariate integrative method to identify reproducible molecular signatures across independent experiments and platforms. BMC Bioinformatics 18, 128. 10.1186/s12859-017-1553-8.

42. Rohart, F., Gautier, B., Singh, A., and Lê Cao, K.-A. (2017). mixOmics: An R package for ‘omics feature selection and multiple data integration. PLoS Comput Biol 13, e1005752. 10.1371/journal.pcbi.1005752.

