## supplementary figures for "Identification of a specific set of genes predicting obesity before phenotype appearance"

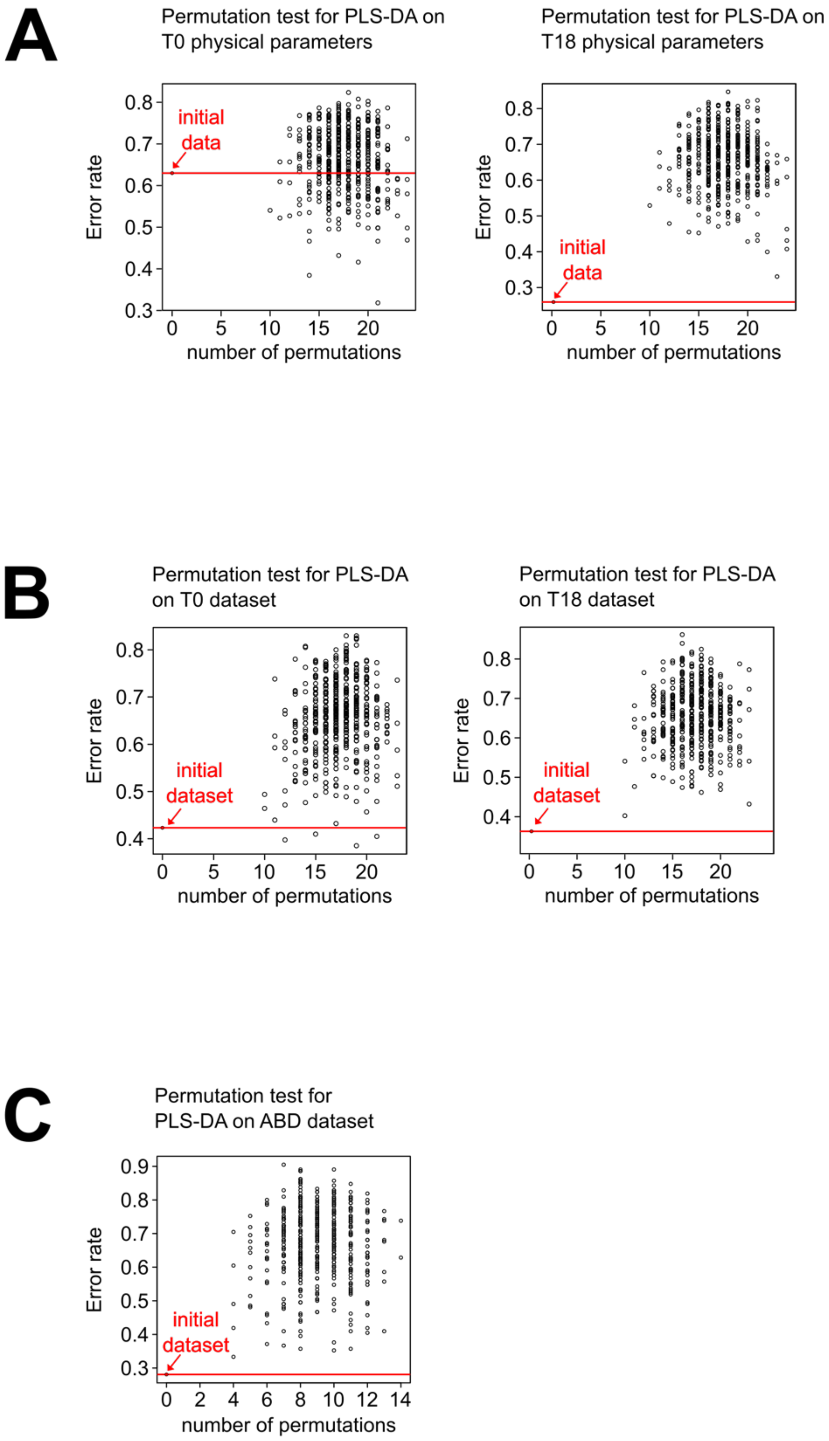
A
Permutation test for PLS-DA on T0 physical parameters
Error rate
0.3
0.4
0.5
0.6
0.7
0.8
initial data
0
5
10
15
20
number of permutations
Permutation test for PLS-DA on T18 physical parameters
Error rate
0.3
0.4
0.5
0.6
0.7
0.8
initial data
0
5
10
15
20
number of permutations
B
Permutation test for PLS-DA on T0 dataset
Error rate
0.4
0.5
0.6
0.7
0.8
initial dataset
0
5
10
15
20
number of permutations
Permutation test for PLS-DA on T18 dataset
Error rate
0.4
0.5
0.6
0.7
0.8
initial dataset
0
5
10
15
20
number of permutations
C
Permutation test for PLS-DA on ABD dataset
Error rate
0.3
0.4
0.5
0.6
0.7
0.8
0.9
initial dataset
0
2
4
6
8
10
12
14
number of permutations

**Supplementary Figure S1:**

The efficiency of the various models obtained was tested using a permutation test. A permutation test is used to assess whether the observed discrimination between groups is statistically significant or could have occurred by chance. The permutation test for PLS-DA involved randomly permuting the group membership of the observations and recalculating the PLS-DA model for each permutation. This procedure was repeated 500 time. The performance of each of the permuted models obtained was assessed by their classification error rate and compared to the error rate of the true model (indicated in red).

**A: Permutation test for PLS-DA with physical parameters measured at T0 and T18.** At T0, the error rates obtained with permuted models were generally as "good" as those obtained with the real dataset (left panel), indicating that it is not possible to create a discriminant model with the T0 physical data. On the other hand, PLS-DA and permutation tests performed on the T18 physical data (i.e. after the HFD-challenge, right panel) confirmed that the three groups identified are indeed different and discrimination can be made based on physical parameters.

**B: Permutation test for PLS-DA with T0 and T18 gene expression datasets**

At both T0 and T18, the error rates obtained with permuted models were higher than the original model showing that the PLS-DA models were valid.

**C: Permutation test for PLS-DA on ABD dataset.**

All error rate values of permuted models were higher than the original model showing that the PLS-DA model was valid.

| | A | B | D |
|---|---|---|---|
| Glucose (mmol/L) | | | |
| Number of values | 26 | 32 | 24 |
| Mean | 8.417 | 7.599 | 8.564 |
| Std. Error of Mean | 0.2441 | 0.1286 | 0.2548 |
| p-value A vs B / A vs D | | 0.013 | ns |
| Cholesterol (mmol/L) | | | |
| Number of values | 15 | 20 | 23 |
| Mean | 2.293 | 2.179 | 2.204 |
| Std. Error of Mean | 0.08365 | 0.08096 | 0.08265 |
| p-value A vs B / A vs D | | ns | ns |
| Triglycerides (mmol/L) | | | |
| Number of values | 14 | 18 | 20 |
| Mean | 0.65 | 0.4917 | 0.809 |
| Std. Error of Mean | 0.02722 | 0.01984 | 0.04611 |
| p-value A vs B / A vs D | | 0.011 | <0.0001 |
| HDL (mmol/L) | | | |
| Number of values | 14 | 20 | 24 |
| Mean | 2.433 | 2.48 | 2.446 |
| Std. Error of Mean | 0.04602 | 0.0683 | 0.0835 |
| p-value A vs B / A vs D | | ns | ns |

**Supplementary Figure S2:**

Plasma metabolites parameters of 5-mo-old male mice from group A, B, and D, were measured in plasma of over-night starved animals.

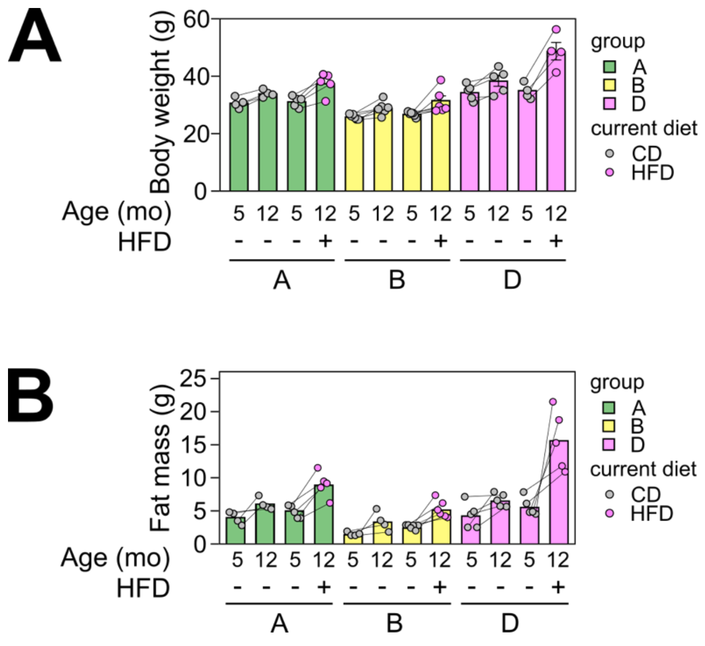


**Supplementary Figure S3:**

**Body weight and fat mass change following a 7-months HFD-challenge on groups A, B and D:** 5-months-old male offspring from groups A, B and D were fed either a CD or a HFD from the age of 5 to 12-month (7 months long HFD-challenge) as described Figure 3C. Body weight and fat mass were recorded at the beginning and at the end of the HFD-challenge. **A: Body weight. B: Fat mass.** n = 4 to 6

**A**

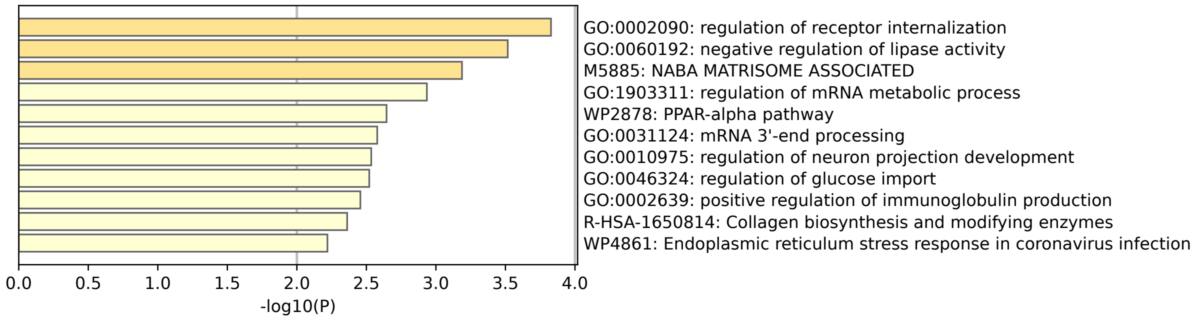


**B**

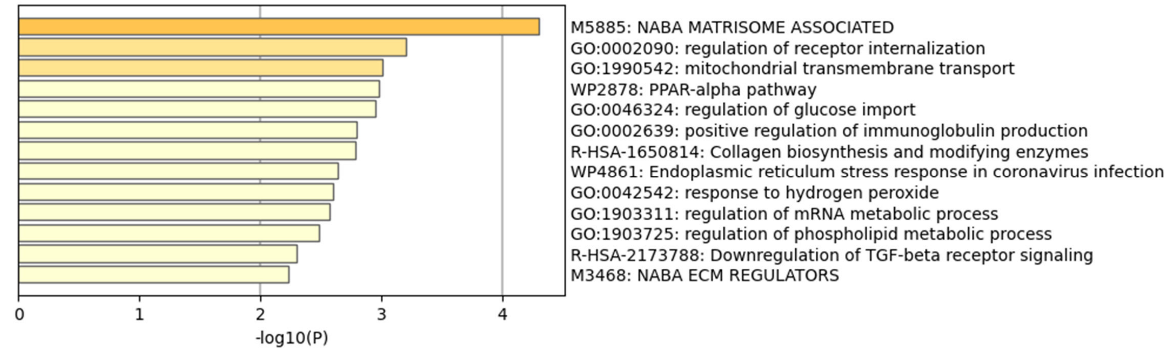


**Supplementary Figure S4: Gene Ontology (GO) enrichment analysis**

The GO enriched terms are colored by p-value[25]. A: enrichment analysis performed on the list of 143 genes (see Table S1, column B). B: enrichment analysis performed on the list of 109 genes (see Table S1, column D)
